# A conserved regulatory switch controls phosphatase activity and specificity

**DOI:** 10.1101/784843

**Authors:** Kristin Ho, Niels Bradshaw

## Abstract

Protein phosphatases must be regulated and specific for their substrates; how this control is achieved is critical for signaling by reversible phosphorylation. We recently reported the discovery of a regulatory switch that controls the activity of the PP2C family serine/threonine phosphatase SpoIIE from the bacterium *Bacillus subtilis*. This regulatory switch activates SpoIIE during spore development by forming the catalytically essential metal-binding site that is conserved across the PP2C family. We hypothesized that this switch is a conserved platform for regulating other PP2C phosphatases. An orthologous phosphatase from *B. subtilis*, RsbU, is activated under stress conditions and responds to different signals and acts on a different phosphoprotein substrate than SpoIIE. Using a combination of biochemical and genetic approaches, we find that broad features of the regulatory mechanism are conserved between SpoIIE and RsbU but that each phosphatase has adapted its response to be most appropriate for the distinct biological outputs it controls. In both cases, the switch accomplishes this by integrating substrate binding and recognition with regulatory inputs to control metal cofactor binding and catalysis. Thus, the switch is a conserved and mechanistically flexible regulatory platform that controls phosphatase activity and substrate specificity.

## Introduction

Protein kinases and phosphatases regulate diverse biological processes by controlling the phosphorylation state of target proteins, requiring that they select specific substrates and act on them at the correct time and place. Protein kinases have conserved allosteric elements that allow control by regulatory inputs and often rely on deep active site grooves to facilitate recognition of cognate substrates (Huse and Kuriyan, 2002; Taylor and Kornev, 2011; Ubersax and Ferrell, 2007). In contrast, the substrate-binding sites of phosphatases are generally surface-exposed and accessible to solvent (Shi, 2009), raising the question: How do protein phosphatases achieve regulation and specificity?

Serine/threonine phosphatases of the PP2C family are precisely regulated to control processes including stress response, development, virulence, and cell growth and death (Lammers and Lavi, 2007; Shi, 2009). Here we focus on two orthologous protein phosphatases, SpoIIE and RsbU, from the bacterium *B. subtilis* that demonstrate how PP2C phosphatases can be subject to distinct regulation and achieve specificity (Fig. 1A). The phosphatases share related enzymatic domains (17% sequence identity, 40% similarity) but have unrelated regulatory domains, respond to different signals, and discriminate between each other’s phosphoprotein substrates.

**Figure 1.**
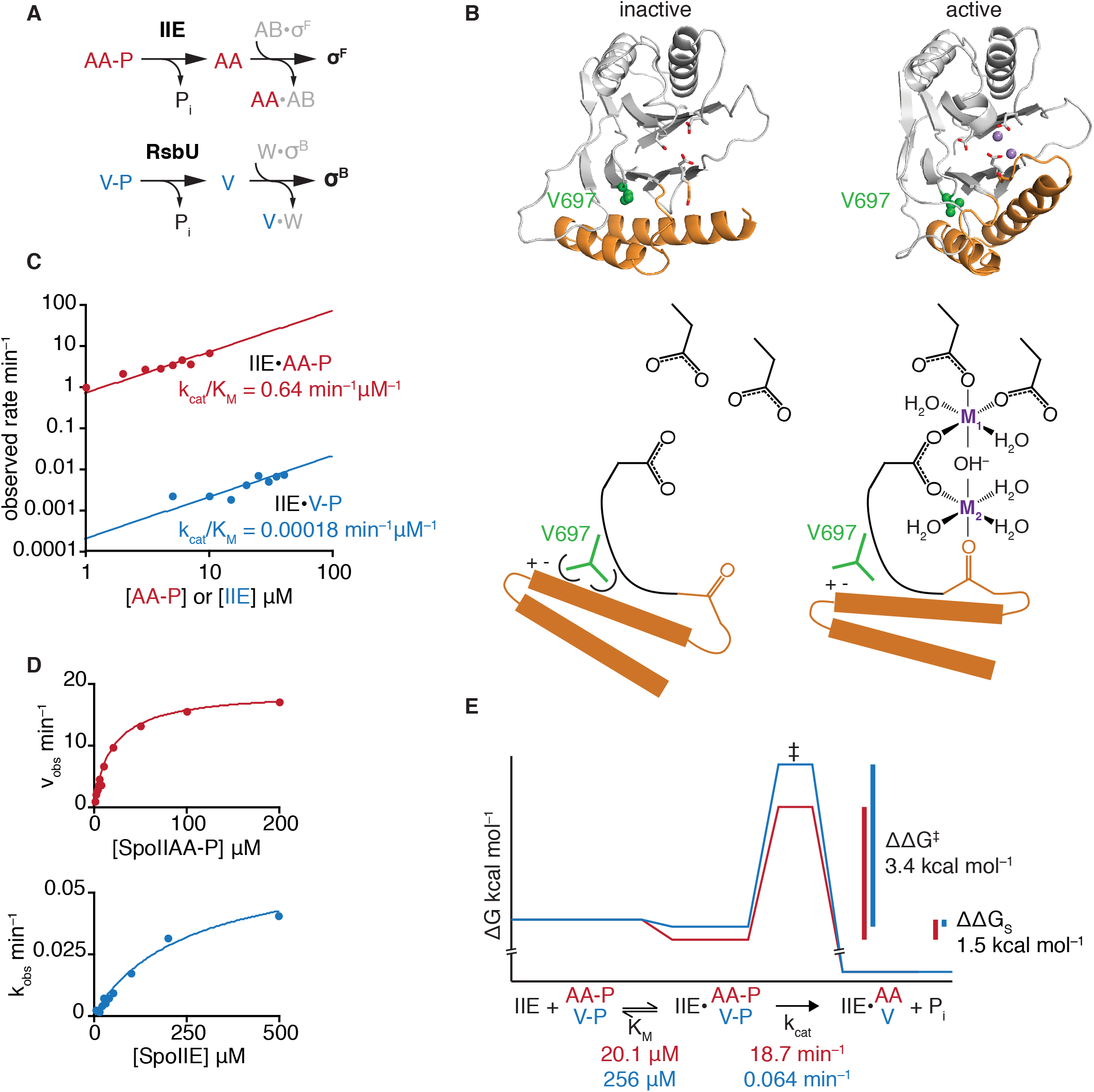
SpoIIE is specific. **A** is a diagram of the pathways controlling activity of the transcription factors σ^F^ and σ^B^. The phosphatases SpoIIE (IIE) and RsbU dephosphorylate paralogous substrate proteins SpoIIAA-P (AA-P) and RsbV-P (V-P). In the unphosphorylated state, SpoIIAA and RsbV bind to anti-sigma-factor proteins SpoIIAB (AB) and RsbW (W), displacing and activating the sigma factors. **B** shows structures of SpoIIE in the inactive (PDBID 5MQH) and active (PDBID 5UCG) states. The α-helical switch is represented in orange and the sidechains that coordinate divalent cations in the active site are shown as sticks. Manganese ions are modeled in the active site of SpoIIE from the structure of metal bound RsbX and are represented as spheres. Valine 697 is shown as green spheres. Below is a cartoon representation of the active site as it is controlled by the switch. **C** is a plot of SpoIIE^590-827^ phosphatase activity with SpoIIAA-P (red) and RsbV-P (blue) as substrate. Reactions with SpoIIAA-P as substrate were performed as multiple turnover reactions with 0.1µM SpoIIE^590-827^ and varying concentrations of SpoIIAA-P. Slow reactions with RsbV-P were performed as single turnover reactions (due to the extremely slow rates of reaction), using trace concentrations of RsbV-P with varying concentrations of SpoIIE^590-827^. All reactions contained 10mM MnCl_2_. Data are fit to the equation observed rate = (k_cat_/K_M_)*[SpoIIAA-P or SpoIIE] and are plotted on log/log axes. **D** is plots of the data from C including higher concentrations of SpoIIAA-P and SpoIIE. Data are fit to the equation observed rate = k_cat_*[SpoIIAA-P or SpoIIE]/(K_M_+ [SpoIIAA-P or SpoIIE]). **E** is a reaction coordinate diagram summarizing the data from **C** and **D**.

SpoIIE dephosphorylates the phosphoprotein SpoIIAA-P to activate a cell-specific transcription factor (σ^F^) during the developmental program of spore formation (Figure 1A) (Duncan et al., 1995; Stragier and Losick, 1996). We recently discovered that SpoIIE is activated by movement of an α-helical element at the base of the phosphatase domain that rotates to form the binding site for the metal cofactor Mn^2+^ (Figure 1B) (Bradshaw et al., 2017). This α-helical switch is a conserved feature of the PP2C phosphatase domain, and we hypothesized that it is a common element for regulating phosphatase activity.

Under conditions of environmental stress, RsbU binds to a partner protein (RsbT) that activates it to dephosphorylate RsbV-P, which, in turn, activates the stress response transcription factor σ^B^ (Figure 1A) (Yang et al., 1996). In addition to sensing different signals, SpoIIE and RsbU exhibit different response profiles; whereas SpoIIE initiates a switch-like developmental transition, RsbU initiates responses with amplitudes graded according to the strength of input (Cabeen et al., 2017; Locke et al., 2011). SpoIIE and RsbU must also be specific for their substrates (Carniol et al., 2004); aberrant dephosporylation of RsbV-P during sporulation blocks spore development (Rothstein et al., 2017) and dephosphorylation of SpoIIAA-P by RsbU would be lethal.

## Results

### SpoIIE principally achieves specificity by stabilizing the transition state

To assess how SpoIIE achieves specificity for its cognate substrate SpoIIAA as opposed to its non-cognate substrate RsbV, we used the C-terminal catalytic domain in isolation (SpoIIE^590-827^) (Fig. 1B). We selected this construct to test the hypothesis that specificity can be encoded by the PP2C catalytic domain alone and to identify potentially conserved mechanisms of specificity. At least three steps of the phosphatase reaction could in principle contribute to specificity: substrate binding (K_M_), catalysis (k_cat_), and cofactor binding. Initially, the metal cofactor concentration was held constant at saturating levels (10 mM MnCl_2_) to isolate substrate binding and catalysis. Under these conditions, SpoIIE^590-827^ was specific, dephosphorylating SpoIIAA-P approximately 3,500-fold more efficiently than the off-pathway substrate, RsbV-P (SpoIIAA-P k_cat_/K_M_ = 0.64 min^-1^µM^-1^, RsbV-P k_cat_/K_M_ = 0.00018 min^-1^µM^-1^) (Fig. 1C). This specificity was principally the result of an increased k_cat_ for the cognate substrate (18.7 min^-1^ for SpoIIAA, 0.064 min^-1^ for RsbV) but was enhanced by a lower K_M_ for the cognate substrate (20 µM for SpoIIAA, 255 µM for RsbV) (Fig.1D). Thus, specificity is achieved by directing 3.4 kcal/mol of the specific binding energy for the cognate substrate to stabilizing the transition state and 1.5 kcal/mol to stabilization of the enzyme substrate complex (Fig 1E).

### The regulatory switch couples substrate binding to cofactor binding

The conserved regulatory switch moves to activate SpoIIE by recruiting metal cofactor to the active site (Fig. 1B). We hypothesized that substrate binding is favored when the switch is in the active conformation, coupling substrate binding to cofactor binding. To investigate this hypothesis, we took advantage of a gain-of-function mutant of SpoIIE (valine 697 to alanine, V697A) in which substitution of a single amino acid in the hydrophobic core of the phosphatase domain biases the switch to the active state (Fig. 1B). The V697A substitution decreased the specificity of SpoIIE, increasing the k_cat_/K_M_ for RsbV-P 148-fold compared to a 5-fold increase for SpoIIAA-P. In the experiments described below, we analyzed the effect of the V697A substitution on K_M_, k_cat_, and cofactor binding with both the cognate and non-cognate substrates.

#### Substrate binding (K_M_)

The V697A substitution decreased the K_M_ of SpoIIE^590-827^ for both SpoIIAA-P (10-fold) and RsbV-P (5-fold) (Fig. 2A), suggesting that substrate binding is favored when SpoIIE is in the active conformation.

**Figure 2.**
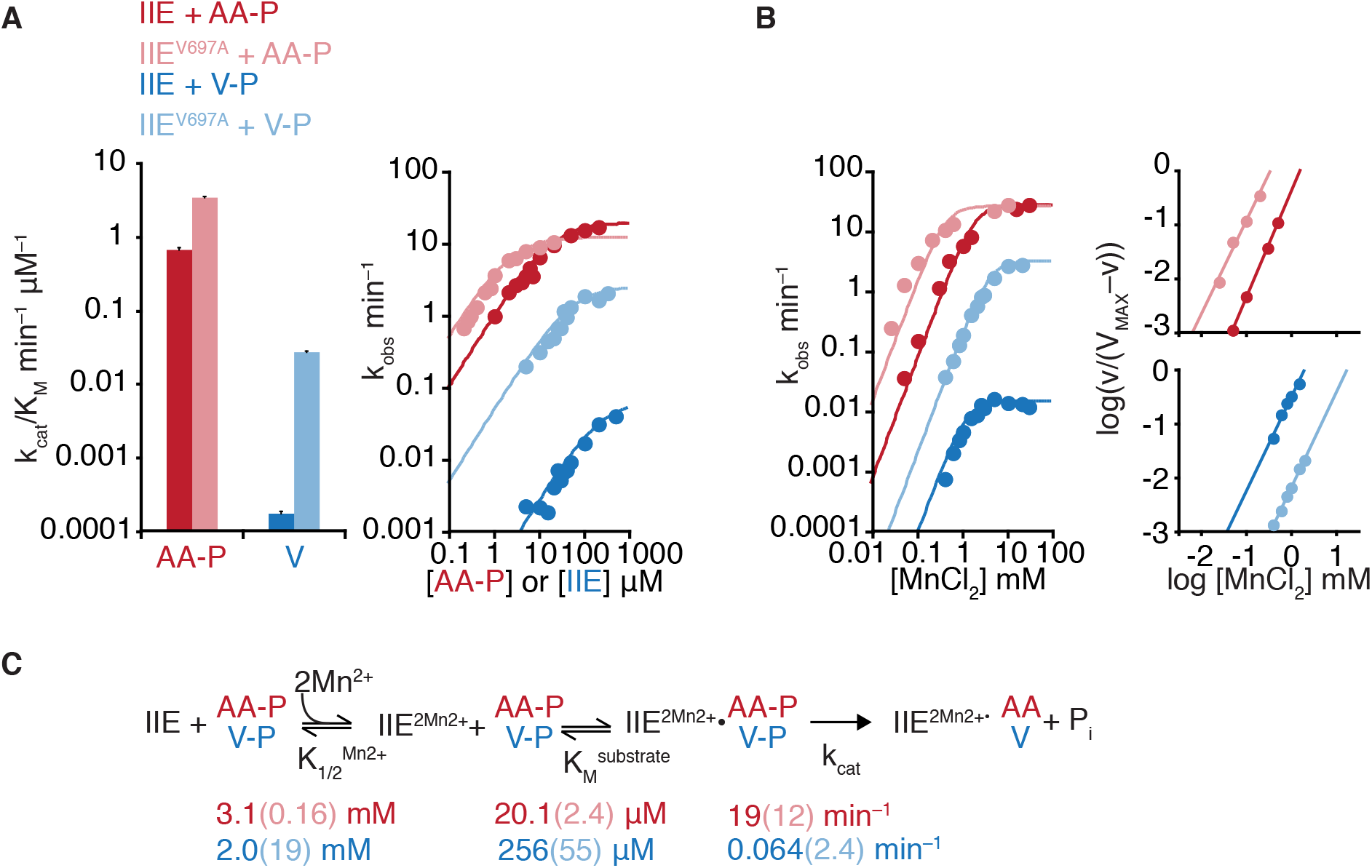
SpoIIE specificity is controlled by the switch. **A** is a bar plot (left) summarizing the impact of the V697A substitution on the k_cat_/K_M_ of SpoIIE^590-827^. Data from SpoIIE^V697A^ are displayed in light colors. Reactions with SpoIIAA-P as substrate were performed as multiple turnover reactions with 0.04µM SpoIIE^590-827 V697A^ and varying concentrations of SpoIIAA-P (this is a lower concentration than used for SpoIIE^590-827^ because of the reduced K_M_ for SpoIIAA-P). Slow reactions with RsbV-P were performed as single turnover reactions (due to the extremely slow rates of reaction), using trace concentrations of RsbV-P with varying concentrations of SpoIIE^590-827 V697A^. All reactions contained 10mM MnCl_2_. To calculate k_cat_/K_M_, data from concentrations below the K_M_ are fit to the equation observed rate = (k_cat_/K_M_)*[SpoIIAA-P or SpoIIE]. Error bars are the error of the fit. Right is a plot of the data including higher concentrations of SpoIIAA-P and SpoIIE. Data are fit to the equation observed rate = k_cat_*[SpoIIAA-P or SpoIIE]/(K_M_+ [SpoIIAA-P or SpoIIE]). Data for wt SpoIIE are from the same experiment as shown in Figure 1D. **B** is a plot of SpoIIE dephosphorylation of SpoIIAA-P (red) and RsbV-P (blue) as a function of MnCl_2_ concentration. Multiple turnover reactions with SpoIIAA-P as substrate included 0.1µM SpoIIE and 50µM SpoIIAA-P. Single turnover reactions with RsbV-P as substrate included 50µM SpoIIE and trace RsbV-P. Data are fit to a cooperative model using the equation k_obs_ = k_cat_*[MnCl_2_]^2^/(K_1/2_+ [MnCl_2_]^2^). Hill plots from the data are shown to the right and are fit to a linear equation. For SpoIIE^590-827^ with SpoIIAA-P *h* is 2.0 (1.7 for V697A) and 1.7 with RsbV-P (1.8 for V697A). **C** is a summary of the reaction scheme for SpoIIE dephosphorylation with values determined for K_1/2_ for MnCl_2_, K_M_ for substrate, and k_cat_ below.

#### Catalysis (k_cat_)

The V697A substitution increased the maximal catalytic rate (k_cat_) of SpoIIE^590-827^ for the non-cognate substrate nearly forty-fold while having a modest effect on the cognate substrate (Fig. 2A). Thus, the switch controls the catalytic rate of SpoIIE and the conformational flexibility of the switch can differentially affect the catalytic rate for cognate and non-cognate substrates.

#### Cofactor binding

To monitor metal cofactor recruitment, we held the substrate concentration constant and measured SpoIIE activity with varied concentrations of MnCl_2_ (Fig. 2B). Consistent with our previous observations, the V697A substitution decreased the concentration of MnCl_2_ required for SpoIIE^590-827^ to dephosphorylate SpoIIAA-P (K_1/2_ 3.1 mM for WT SpoIIE^590-827^ and K_1/2_ 0.16 mM for SpoIIE^590-827^ V697A). In contrast, the V697A substitution increases the MnCl_2_ concentration required for dephosphorylation for the non-cognate substrate RsbV-P (K_1/2_ of 2.0 mM for WT SpoIIE^590-827^and K_1/2_ of 19 mM for SpoIIE^590-827^ V697A). This suggests that the switch mediates coupling between cofactor binding, substrate binding, and catalysis and that the V697A substitution shifts binding energy for the non-cognate substrate from cofactor binding to catalysis (summarized in Fig. 2C). Thus, at concentrations of MnCl_2_ that are below the K_1/2_, SpoIIE^590-827 V697A^ retains specificity (approximately a thousand-fold less active towards RsbV-P than SpoIIAA-P), but specificity is reduced at saturating cofactor concentrations. Cooperativity of metal cofactor binding (Hill coefficient of two, inset panels Fig. 2B), yields a switch-like response profile, appropriate for initiation of a developmental transition.

### The SpoIIE regulatory domain controls cofactor binding, substrate binding and catalysis

PP2C phosphatases have regulatory domains that control their activity(Bradshaw et al., 2017; Shi, 2009). We therefore repeated the above analysis with the phosphatase domain of SpoIIE together with a portion of the regulatory domain (SpoIIE^457-827^). We chose this fragment because it is a well-behaved soluble protein whose crystal structure we previously reported (Bradshaw et al., 2017) (Figure 2 – figure supplement 1). Although the regulatory domain does not receive a native signal with this truncated protein, the combined phosphatase and regulatory domains represent a simple system for assessing how accessory domains can interface with the switch. SpoIIE^457-827^ was specific for SpoIIAA-P (SpoIIAA-P k_cat_/K_M_ = 0.039 min^-1^µM^-1^, RsbV-P k_cat_/K_M_ = 7.2 x10^-5^ min^-1^µM^-1^) (Figure 2 – figure supplement 1A). Additionally, the V697A substitution reduces the selectivity for SpoIIAA-P; increasing the activity towards RsbV-P 100-fold, while not affecting activity towards SpoIIAA-P (Figure 2 – figure supplement 1A).

However, SpoIIE^457-590^ exhibited several notable differences from the phosphatase domain alone (SpoIIE^590-827^). First, the regulatory domain increased the K_M_ of SpoIIE for both cognate and non-cognate substrates such that the concentration dependence of phosphatase activity was linear even at concentrations of substrate above 100 µM (Figure 2 – figure supplement 1A). Second, the regulatory domain altered how the V697A substitution affected cofactor binding; whereas the substitution decreased the concentration of MnCl_2_ required for SpoIIE^457-827^ to dephosphorylate SpoIIAA-P (Figure 2A), it had no effect on the concentration of MnCl_2_ required to dephosphorylate RsbV-P (Figure 2 – figure supplement 1B).

Together these findings (summarized in Figure 2 – figure supplement 1C) are consistent with coupling between substrate binding, catalysis, and cofactor recruitment and indicate that regulatory domains can influence all three steps through the switch.

### Mechanisms of regulation and specificity are conserved between SpoIIE and RsbU

To test whether the α-helical switch is a conserved element for PP2C phosphatase regulation and specificity, we examined RsbU, the cognate phosphatase for RsbV-P. We hypothesized that RsbT activates RsbU by stimulating cofactor binding through the switch. Consistent with a role for the switch, we found that RsbT decreased the K_M_ of RsbU for Mn^2+^ by nearly 100-fold (K_M_ 0.98 µM in the presence of RsbT and 77 µM in the absence of RsbT) (Fig. 3B). In contrast to SpoIIE, RsbU activity was non-cooperative with respect to Mn^2+^ concentration (Fig. 3B right panels). Whereas SpoIIE regulates a developmental switch, and thus requires an all-or-nothing response, RsbU regulates a stress response, for which a graded response may be advantageous.

**Figure 3.**
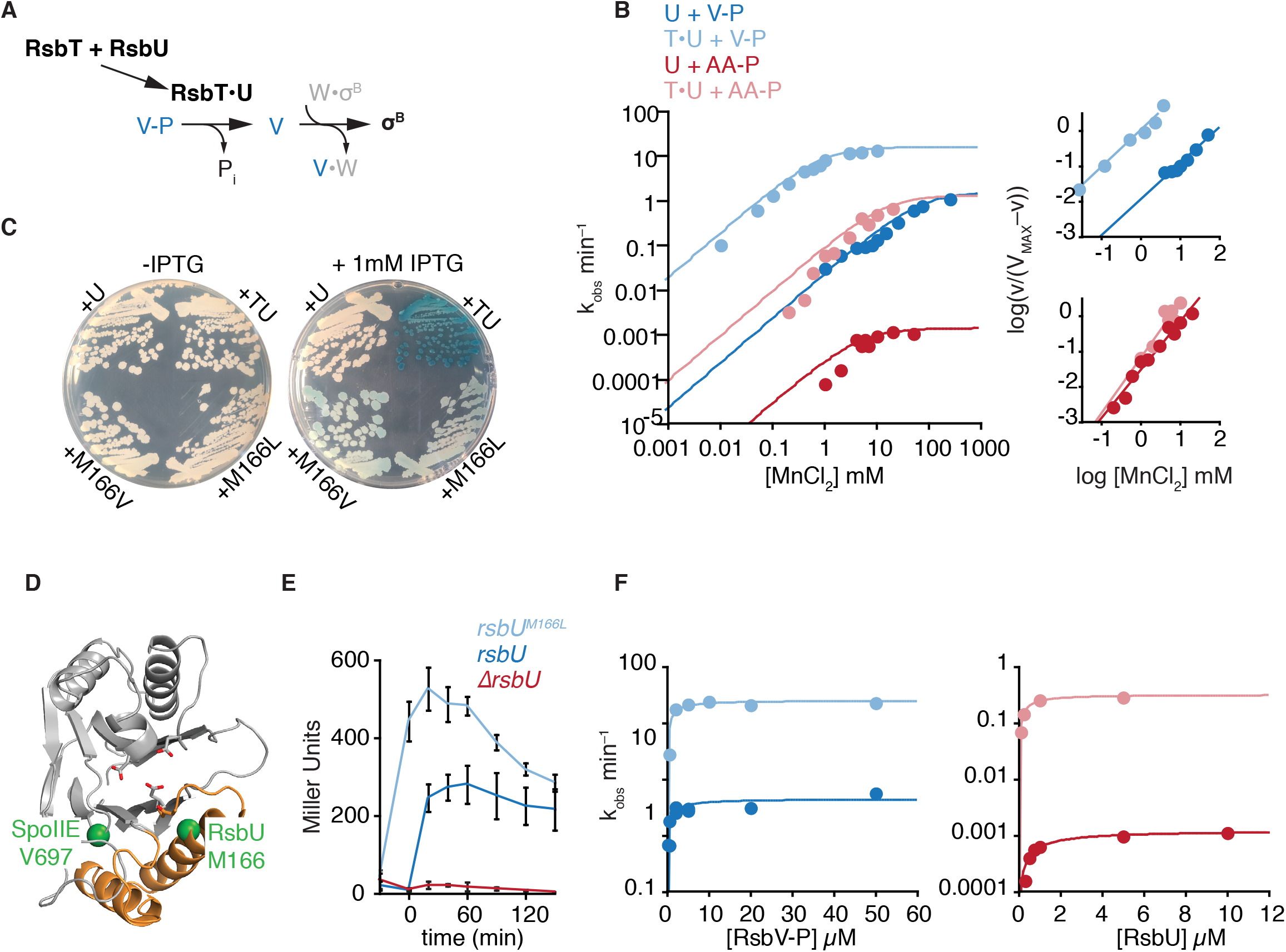
The switch controls RsbU activity and specificity. **A** is a schematic of RsU activation by RsbT to activate σ^B^ in response to stress. **B** is a plot of RsbU dephosphorylation of RsbV-P (blue) and SpoIIAA-P (red) in the presence (light color) and absence (dark color) of 10µM RsbT. Reactions were performed at varying concentrations of MnCl_2_ and data are fit to the equation k_obs_ = k_cat_*[MnCl_2_]/(K_M_+ [MnCl_2_]). To the right are Hill plots including data from 10% to 90% activity fit to a linear equation. For RsbU with RsbV-P *h* is 1.0 (1.0 for +T) and 1.4 with SpoIIAA-P (1.6 for +T). For reactions with SpoIIAA as substrate, single turnover reactions with 2.5µM RsbU were performed (due to the slow rates of dephosphorylation). Reactions with RsbV-P as substrate included 25µM RsbU (single turnover in the absence of T) or 25µM RsbV-P and 0.5µM RsbU (multiple turnover in the presence of T because of the rapid rate of dephosphorylation). **C** is the result of a blue-white screen of a stress non-responsive strain carrying a reporter for SigB activity (Δ*rsbPQ*Δ*rsbTU rsbV-flag amyE::ctc-lacZ*) carrying plasmid containing *rsbU* variants or *rsbT* and *rsbU* under the control of an IPTG-inducible reporter. Starting from the upper left and moving clockwise: *rsbU, rsbTU, rsbU^M166L^*, *rsbU^M166V^*. Strains were grown on LB plates containing MLS (for plasmid retention), 80 µg/mL X-gal, and either 0 or 1 mM IPTG for 2 days at 37°C. Presence of blue pigment indicates SigB activity and activation of the stress response pathway. Expression of *rsbTU, rsbU^M166V^,* or *rsbU^M166L^,* but not wild-type *rsbU*, activates the stress response system. **D** is a structure of the SpoIIE PP2C domain (from PDBID 5UCG). The α-helical switch is represented in orange and the sidechains that coordinate divalent cations in the active site are shown as sticks. The positions of SpoIIE, Valine 697 and RsbU Methionine 166 are marked with green spheres. **E** is the result of a beta-galactosidase assay of ethanol-stressed *rsbPQ* deleted strains carrying a reporter for σ^B^ activity (Δ*rsbPQ amyE::ctc-*lacZ) either encoding wild-type *rsbU* (dark blue) or *rsbU^M166L^* (light blue) on the chromosome, or deleted for *rsbU* (red). Cultures were grown in LB medium to late-log phase and then stressed with addition of 4% ethanol (final concentration). Cultures were sampled before (t = −30, 0) and after stress induction and *lacZ* expression was analyzed using ONPG as a substrate. Values plotted are the means of three independent replicates, and the error bars are standard deviations of the mean. **F** is plots of RsbU activity towards RsbV-P (left, blue) and SpoIIAA-P (right, red) in the absence (dark colors) and presence (light colors) of RsbT (10µM). Curves are fits to the Michaelis-Menton equation rate = k_cat_*[RsbU or RsbV-P]/(K_M_+ [RsbU or RsbV-P]). The k_cat_ for RsbV-P is 25 min^−1^ (0.4 min^−1^ without T) and K_M_ 1.3 µM (1.5 µM without T). The k_cat_ for SpoIIAA-P is 0.32 min^−1^ (0.0012 min^−1^ without T) and K_M_ 0.3 µM (1 µM without T). Multiple turnover reactions included 0.1µM RsbU and all reactions were performed with saturating MnCl_2_ (100mM without T or 10mM in the presence of T).

In our *in vitro* assays, RsbT additionally enhanced the maximal catalytic activity of RsbU towards RsbV-P 10-fold (k_cat_ 15 min^−1^ in the presence of RsbT and 1.5 min^−1^ in the absence of RsbT) (Fig. 3B), consistent with previous observations (Delumeau et al., 2004; Yang et al., 1996). This suggests that, as we observed for SpoIIE, control of catalysis and cofactor binding are coupled, providing nearly a 1000-fold increase in the activity of RsbU at physiological concentrations of metal ion. The magnitude of effects of RsbT are lower limits because RsbT solubility restricted how much RsbT could be added to reactions.

The above observations reveal striking similarities to how the switch controls SpoIIE activity and cofactor-binding, suggesting a role for the switch in RsbU activation. To assess this possibility, we took an unbiased approach, performing a genetic screen to isolate variants of RsbU that are active in the absence of RsbT. In a SigB-reporter strain deleted for both pathways known to dephosphorylate RsbV-P (Δ*rsbPQ* Δ*rsbTU rsbV-FLAG amyE::ctc-lacZ*), we expressed a library of randomly mutagenized RsbU under control of an IPTG inducible promoter. Using lacZ production from the σ^B^ controlled promoter, we identified two amino acid substitutions in the phosphatase domain, both at M166 (M166V, and M166L), that activated σ^B^ in the absence of RsbT (Figure 3C). Signal from the σ^B^ dependent LacZ reporter appeared two days after induction of RsbU M166L (in contrast to induction of RsbTU, which appeared more intensely and after 24 hours), suggesting that substitution of M166 is a partial bypass of RsbT activation. M166 is in the N-terminal α-helix of the switch and is predicted to pack in the hydrophobic core of the phosphatase domain (Figure 3D).

We hypothesize that substitution of M166 with valine or leucine favors the active state of the RsbU switch, analogous to V697 substitution of SpoIIE. One prediction of this hypothesis is that RsbU^M166L^ would elevate basal σ^B^ activity that is enhanced by RsbT. To test this, we analyzed σ^B^ activity in strains with *rsbU* variants at the native locus that retain *rsbT* but lack *rsbPQ* (the RsbTU-independent branch of the general stress response pathway). Samples were taken from mid-log phase cultures 30 minutes before stress-induction (4% ethanol), immediately before stress-induction, and then at regular intervals (Figure 3E). For both the wild-type *rsbU* and *rsbU^M166L^* strains, σ^Β^ activity peaked after stress induction and then decreased, suggesting that RsbU^M166L^ is responsive to RsbT. Νο σ^B^ activity was observed in a strain lacking *rsbU*. There were two major differences caused by the M166L substitution. First, the *rsbU^M166L^* strain had overall higher levels of σ^Β^ activity. Second, the basal level of σ^Β^ activity observed for the *rsbU^M166L^* encoding strain increased sharply as the cells reached late-log phase but before the stress induction (t = 0). Indeed, this pre-stress activity exceeded the peak level of σ^Β^ activity observed for the wild-type *rsbU* strain. RsbU is known to be activated when cells enter stationary phase suggesting that the M166L substitution potentiates RsbU. Together with the results from the genetic screen, this suggests that RsbU^M166L^ is hyperactivated in the presence and absence of RsbT and indicates a central role for the regulatory switch in controlling RsbU phosphatase activity.

### RsbU is Specific

Reconstitution of RsbU activation by RsbT further allowed us to determine how specificity is maintained upon phosphatase activation. Like SpoIIE, RsbU is specific: RsbU had higher catalytic activity towards its cognate substrate (k_cat_ 0.4 min^−1^ for RsbV-P and 0.0012 min^− 1^ for SpoIIAA-P), and bound both cognate and non-cognate substrates tightly (K_M_ 1.1µM for SpoIIAA-P and 1.5 µM for RsbV-P; K_M_ values were difficult to determine accurately because of limitations of detection for substrate proteins) (Fig. 3F).

As with the cognate substrate, RsbT enhanced RsbU catalytic activity towards SpoIIAA-P (a 300-fold increase to k_cat_ 0.32 min^−1^) (Fig. 3F). However, similar to the V697A mutation of SpoIIE, RsbT did not substantially change the affinity of cofactor binding with the non-cognate substrate(K_M_ 5.5 mM in the absence of RsbT and 16 mM in the presence of RsbT) (Fig. 3B). RsbT also had little impact on SpoIIAA-P binding (approximate K_M_ 1 µM in the absence of RsbT and 0.3 µM in the presence of RsbT) (Fig. 3F). Thus, specificity is maintained because RsbT activation fails to stimulate cofactor binding for RsbU in the presence of its non-cognate substrate.

## Discussion

By examining two phosphatases that are regulated by different signals and that discriminate between each other’s substrate proteins, we found that the mechanisms of PP2C regulation and specificity are structurally conserved but functionally flexible to produce different response profiles for different systems. This is achieved by a regulatory switch that integrates input signals and binding to cognate substrate to control cofactor binding in the active site and catalytic activity. Although allosteric regulatory mechanisms don’t generally directly provide substrate specificity (Herschlag, 1998), an allosteric regulatory mechanism that couples regulatory inputs, catalysis, and substrate recognition provides a mechanism to control diverse signaling pathways that is evolutionarily adaptable.

This discovery raises the question: How does the switch coordinate regulatory inputs, substrate binding, cofactor binding, and catalysis? A possible simple mechanism is that both regulatory domains and substrates directly contact and influence the conformation of the switch. Additional control of the maximal catalytic rate could be achieved based on how the orientation of the switch determines the position of the phosphoserine bond relative to the active site or the geometry of the metal cofactors. This seems plausible because the switch is positioned just below the active site where it would likely contact substrate proteins. In the single available structure of a PP2C phosphatase bound to its substrate protein (Hab1 bound to SnRK2), SnRK2 makes direct contacts with the Hab1 switch and to a variable region (termed “the flap”) that packs against the switch (Soon et al., 2012). Regulatory domains from diverse PP2C phosphatases similarly pack against the switch (Bradshaw et al., 2017; Dorich et al., 2019; Vassylyev and Symersky, 2007; Zhang et al., 2013). For different phosphatases, this coupling could be used to recruit substrate when the phosphatase is in its active conformation or to ensure that the phosphatase only adopts the active conformation when bound to the cognate substrate.

## Materials and Methods

### Growth conditions

Liquid cultures were grown in Lennox lysogeny broth (LB) while colonies were grown on LB containing Bacto agar. The antibiotics used were MLS (50 µg/mL erythromycin, 250 µg/mL lincomycin), chloramphenicol (20 µg/mL for *E. coli*), carbenicillin (100 µg/mL), and kanamycin (10 µg/mL for *B. subtilis* or 50 µg/mL for *E. coli*). When indicated, 80 µg/mL of 5-bromo-4-chloro-3-indolyl-β-D-galactopyranoside (X-gal) and 1 mM Isopropyl β-d-1-thiogalactopyranoside (IPTG) were added to growth medium to visualize SigB reporter activity on plates.

### Protein expression constructs

SpoIIAA and SpoIIAB were expressed in BL21(DE3) Rosetta2 pLysS cells as 6H-sumo fusions in pET23a (made by ligation of the coding sequence in pET23a 6H-sumo digested with NotI/AgeI as reported (Bradshaw and Losick, 2015)). SpoIIAA-P was produced in BL21(DE3) cells containing pET-YSBLIC 6H-3C-spoIIAA–spoIIAB as reported (Levdikov et al., 2012). SpoIIE^590-827^, SpoIIE^457-827^, RsbT, RsbU, RsbV, and RsbW were produced with 3C cleavable 6H tags in BL21(DE3) cells (constructs were made by insertion of the coding sequence to pET47b vectors digested with XmaI/XhoI by isothermal assembly). Mutations were introduced to SpoIIE and RsbU expression constructs using QuikChange site directed mutagenesis. RsbV-P was produced by co-expressing RsbV and RsbW using a pET47b vector generated by isolation of the coding sequences for RsbV and RsbW from genomic DNA, yielding a construct with RsbV fused to a 3C cleavable 6H tag followed by untagged RsbW. All strains used are listed in the table of strains.

### Protein expression and purification

Proteins were expressed and purified using slight modifications to previous methods(Bradshaw et al., 2017). All proteins were purified to greater-than 95% purity as assessed by Coomassie stained SDS-PAGE gel. Unless otherwise noted, protein expression was induced with 1mM IPTG for 14-18 hours at 14°C in cells that had been grown to OD_600_ of 0.6 at room temperature. Details for purification of individual proteins follows:

*SpoIIE* (same protocol for all variants) was purified from cell pellets lysed (using a microfluidizer) in 50mM K•HEPES pH 8.0, 200mM NaCl, 10% glycerol, 0.5mM DTT, 20mM imidazole, 0.1mg/mL PMSF. Lysates were clarified by spinning for 30 min at 16,000 RPM in a Sorvall SS-34 rotor at 4°C. Protein was bound to a HisTrap HP column on an AKTA FPLC, was washed, and then was eluted with a gradient to 200mM imidazole. The 6H tag was cleaved overnight in dialysis with PreScission protease, and the tag and protease were removed by flowing over Ni-NTA resin. SpoIIE was next purified on a Resource Q column equilibrated in 50mM HEPES pH8.0, 100mM NaCl, 2mM EDTA, 2mM DTT and eluted with a gradient to 500mM NaCl. The SpoIIE containing fractions were spin concentrated and gel-filtered on a 120ml Superdex 75 column equilibrated in 20mM K•HEPES pH 8.0, 100mM NaCl, 10% glycerol, 2mM DTT. Protein was spin-concentrated to approximately 400µM and flash-frozen and stored at –80°C.

*SpoIIAA* was expressed for 4 hours at 37°C. Cell pellets were resuspended in 20mM K•HEPES pH 7.5, 200mM NaCl, 0.5mM DTT, 0.1mg/ml PMSF (5ml/L of culture). Cells were lysed using a cell disruptor in one-shot mode (Constant Systems, Daventry, United Kingdom) and lysates were clarified by spinning for 30 min at 16,000 RPM in a Sorvall SS-34 rotor at 4°C. Protein was bound to Ni-NTA resin (1 ml/L of culture) on column by gravity flow, washed with buffer containing 20mM imidazole, and eluted with 200mM imidazole. The 6H-sumo tag was cleaved using 10µl of a 100µM stock of ULP1 in overnight dialysis at 16°C to 20mM K•HEPES pH 7.5, 200mM NaCl, 10% glycerol, 0.5mM DTT. The 6H-sumo tag was subtracted by flowing over Ni-NTA resin equilibrated in the cleavage and dialysis buffer. The cleaved protein was then spin-concentrated and gel-filtered on a 120ml Superdex 75 column equilibrated in 50mM K•HEPES pH 8.0, 100mM NaCl, 10% glycerol, 2mM DTT. Protein was concentrated to approximately 400µM, flash-frozen and stored at −80°C.

*SpoIIAB* was expressed for 4 hours at 37°C. Cell pellets were resuspended in 50mM K•HEPES pH 7.5, 200mM NaCl, 10mM MgCl_2_, 0.5% Triton X-100, 0.5mM DTT, 0.1mg/ml PMSF (5ml/L of culture). Cells were lysed using a cell disruptor in one-shot mode (Constant Systems, Daventry, United Kingdom) and lysates were clarified by spinning for 30 min at 16,000 RPM in a Sorvall SS-34 rotor at 4°C. Protein was bound to Ni-NTA resin (1 ml/L of culture) on column by gravity flow, washed with buffer containing 20mM imidazole, and eluted with 200mM imidazole. The 6H-sumo tag was left un-cleaved to aid in removal from phosphorylation reactions. SpoIIAB was spin-concentrated and gel-filtered on a 120ml Superdex 75 column equilibrated in 50mM K•HEPES pH 7.5, 175mM NaCl, 10mM MgCl_2_, 10% glycerol, 2mM DTT. Protein was concentrated to approximately 400µM, supplemented to 50% glycerol and flash-frozen and stored at −80°C.

*SpoIIAA-P* was purified from cells co-expressing 6H-3C-SpoIIAA and untagged SpoIIAB. for 4 hours at 37°C. Cell pellets were resuspended in 20mM K•HEPES pH 7.5, 200mM NaCl, 0.5mM DTT, 0.1mg/ml PMSF (5ml/L of culture). Cells were lysed using a cell disruptor in one-shot mode (Constant Systems, Daventry, United Kingdom) and lysates were clarified by spinning for 30 min at 16,000 RPM in a Sorvall SS-34 rotor at 4°C. Protein was bound to a HisTrap HP column on an AKTA FPLC, was washed with buffer containing 20mM imidazole and 500mM NaCl, and eluted with a gradient to 300mM imidazole. The tag was cleaved using PreScission protease overnight dialysis at 4°C. The cleaved protein was then spin-concentrated and gel-filtered on a 120ml Superdex 75 column equilibrated in 50mM K•HEPES pH 8.0, 100mM NaCl, 10% glycerol, 2mM DTT. Protein was concentrated to approximately 400µM, flash-frozen and stored at –80°C.

*SpoIIAB* was expressed for 4 hours at 37°C. Cell pellets were resuspended in 50mM K•HEPES pH 7.5, 200mM NaCl, 10mM MgCl_2_, 0.5% Triton X-100, 0.5mM DTT, 0.1mg/ml PMSF (5ml/L of culture). Cells were lysed using a cell disruptor in one-shot mode (Constant Systems, Daventry, United Kingdom) and lysates were clarified by spinning for 30 min at 16,000 RPM in a Sorvall SS-34 rotor at 4°C. Protein was bound to Ni-NTA resin (1 ml/L of culture) on column by gravity flow, washed with buffer containing 20mM imidazole, and eluted with 200mM imidazole. The 6H-sumo tag was left un-cleaved to aid in removal from phosphorylation reactions. SpoIIAB was spin-concentrated and gel-filtered on a 120ml Superdex 75 column equilibrated in 50mM K•HEPES pH 7.5, 175mM NaCl, 10mM MgCl_2_, 10% glycerol, 2mM DTT. Protein was concentrated to approximately 400µM, supplemented to 50% glycerol and flash-frozen and stored at –80°C.

*RsbT* was purified from cell pellets lysed (using a microfluidizer) in 20mM K•HEPES pH 7.5, 200mM NaCl, 10% glycerol, 0.5mM DTT, 20mM imidazole, 0.1mg/mL PMSF. Lysates were clarified by spinning for 30 min at 16,000 RPM in a Sorvall SS-34 rotor at 4°C. Protein was bound to Ni-NTA resin (1 ml/L of culture) on column by gravity flow, washed, and eluted with 200mM imidazole. The 6H tag was cleaved overnight in dialysis with PreScission protease, and the tag and protease were removed by flowing over Ni-NTA resin. RsbT was spin concentrated and gel-filtered on a 120ml Superdex 75 column equilibrated in 20mM K•HEPES pH 7.5, 150mM NaCl, 10% glycerol, 2mM DTT. Protein was spin-concentrated to approximately 40µM and flash-frozen and stored at –80°C.

*RsbU* was purified from cell pellets lysed (using a microfluidizer) in 50mM K•HEPES pH 8.0, 200mM NaCl, 10% glycerol, 0.5mM DTT, 20mM imidazole, 0.1mg/mL PMSF. Lysates were clarified by spinning for 30 min at 16,000 RPM in a Sorvall SS-34 rotor at 4°C. Protein was bound to a HisTrap HP column on an AKTA FPLC, was washed, and then was eluted with a gradient to 200mM imidazole. The 6H tag was cleaved overnight in dialysis with PreScission protease, and the tag and protease were removed by flowing over Ni-NTA resin. Cleaved RsbU was spin concentrated and gel-filtered on a 120ml Superdex 75 column equilibrated in 20mM K•HEPES pH 8.0, 100mM NaCl, 10% glycerol, 2mM DTT. The RsbU containing fractions were then purified on a Resource Q column equilibrated in 50mM HEPES pH8.0, 100mM NaCl, 2mM EDTA, 2mM DTT and eluted with a gradient to 500mM NaCl. This final step removes a degradation product present in some RsbU preparations. Protein was supplemented with 10% glycerol, spin-concentrated to approximately 400µM and flash-frozen and stored at –80°C.

*RsbV* was purified from cell pellets lysed (using a microfluidizer) in 20mM K•HEPES pH 7.5, 200mM NaCl, 10% glycerol, 0.5mM DTT, 20mM imidazole, 0.1mg/mL PMSF. Lysates were clarified by spinning for 30 min at 16,000 RPM in a Sorvall SS-34 rotor at 4°C. Protein was bound to Ni-NTA resin (1 ml/L of culture) on column by gravity flow, washed, and eluted with 200mM imidazole. The 6H tag was cleaved overnight in dialysis with PreScission protease, and the tag and protease were removed by flowing over Ni-NTA resin. RsbT was spin concentrated and gel-filtered on a 120ml Superdex 75 column equilibrated in 20mM K•HEPES pH 7.5, 150mM NaCl, 10% glycerol, 2mM DTT. Protein was spin-concentrated to approximately 400µM and flash-frozen and stored at –80°C.

*RsbW* was purified from cell pellets lysed (using a microfluidizer) in 20mM K•HEPES pH 7.5, 10mM MgCl_2_, 200mM NaCl, 10% glycerol, 0.5mM DTT, 20mM imidazole, 0.1mg/mL PMSF. Lysates were clarified by spinning for 30 min at 16,000 RPM in a Sorvall SS-34 rotor at 4°C. Protein was bound to a HisTrap HP column on an AKTA FPLC, washed, and eluted with a gradient to 250mM imidazole. The 6H tag was left un-cleaved to aid removal after phosphorylation reactions. RsbW was then purified on a Resource Q column equilibrated in 50mM HEPES pH 7.5, 200mM NaCl, 10 mM MgCl_2_, 2mM DTT and eluted with a gradient to 1M NaCl. RsbW containing fractions were spin concentrated and gel-filtered on a 120ml Superdex 75 column equilibrated in 50mM K•HEPES pH 7.5, 150mM NaCl, 10% glycerol, 2mM DTT. Protein was spin-concentrated to approximately 100µM and flash-frozen and stored at –80°C.

*RsbV-P* was purified from cells co-expressing 6H-3C-RsbV and untagged RsbW that were grown at 37°C and induced with 1mM IPTG for 4 hours at 37°C. Cells were lysed (using a microfluidizer) in 50mM K•HEPES pH 7.5, 50mM KCl, 10% glycerol, 0.5mM DTT, 20mM imidazole, 0.1mg/mL PMSF. Lysates were clarified by spinning for 30 min at 16,000 RPM in a Sorvall SS-34 rotor at 4°C. Protein was bound to a HisTrap HP column on an AKTA FPLC, washed, and eluted with a gradient to 250mM imidazole. RsbVW containing fractions were then gel-filtered using a 120 ml Superdex 75 column and fractions containing either RsbV or RsbVW complex were collected. The sample was supplemented with 5mM ATP, 10mM MgCl2, and PreScission protease and placed in dialysis in 200ml of buffer also supplemented with MgCl_2_ and ATP at room temperature overnight. The 6H tag and protease were removed by flowing over Ni-NTA resin and the protein was gel-filtered again on a Superdex 75 column equilibrated in 50mM K•HEPES pH 8.0, 100mM NaCl, 2mM DTT, 10% glycerol. Free RsbV-P was collected and complete phosphorylation was confirmed by isoelectric focusing. Protein was spin-concentrated to approximately 400µM and flash-frozen and stored at –80°C.

### Phosphatase Assays

Phosphatase assays were performed using methods reported previously (Bradshaw and Losick, 2015) with modifications as described. To produce ^32^P labeled SpoIIAA-P, 75 µM SpoIIAA, 5 µM SpoIIAB and 50 µCi of γ-^32^P ATP were incubated overnight at room temperature in 50 mM K•HEPES pH 7.5, 50 mM KCl, 750 µM MgCl_2_, 2 mM DTT. Unincorporated nucleotide was removed from the reaction mixture by buffer exchange using a Zeba spin column (Pierce) equilibrated in 20 mM K•HEPES pH 7.5, 200 mM NaCl, 2 mM DTT. The sample was then flowed over Q-sepharose resin to remove SpoIIAB. SpoIIAA-P was then buffer exchanged to the buffer used for subsequent assays using a Zeba spin column equilibrated in 50mM K•HEPES pH 8.0, 100mM NaCl. Labeled SpoIIAA-P was aliquoted and frozen at –80°C for future use.

To produce ^32^P labeled RsbV-P, 50 µM RsbV, 5 µM RsbW and 200 µCi of γ-^32^P ATP were incubated overnight at room temperature in 50 mM K•HEPES pH 7.5, 50 mM KCl, 10 mM MgCl_2_, 2 mM DTT. Unincorporated nucleotide was removed from the reaction mixture by buffer exchange using a Zeba spin column (Pierce) equilibrated in 50 mM K•HEPES pH 8.0, 100 mM NaCl, 20mM imidazole. 6H-tagged RsbW was then removed by binding to Ni-NTA resin. The flowthrough containing RsbV-P was then exchanged to the buffer used for subsequent assays using a two successive Zeba spin columns equilibrated in 50mM K•HEPES pH 8.0, 100mM NaCl to remove all unincorporated nucleotide and free phosphate. Labeled RsbV-P was aliquoted and frozen at –80°C for future use.

All phosphatase assays were performed at room temperature in 25 mM K•HEPES pH 8, 100 mM NaCl, 100 µg/ml BSA (to prevent protein sticking to tubes). The concentrations of enzyme, substrate, and MnCl_2_ were varied as indicated. Reactions were stopped in 0.5M EDTA pH 8.0, 2% Triton X-100 and run on PEI-Cellulose TLC plates developed in 1 M LiCl_2_, 0.8 M Acetic Acid, and imaged on a Typhoon (GE Life Sciences). Phosphatase assays were performed more than three independent times as separate experiments.

### Strain construction

*B. subtilis* strains were constructed using standard molecular genetic techniques (Harwood and Cutting, 2010) in the PY79 strain background (Youngman et al., 1984; Zeigler et al., 2008) and were validated to contain the correct constructs by double-crossover recombination at the correct insertion site. All strains are used in this study are described in table of strains (Table 1). A SigB reporter strain (KC479) previously described (Carniol et al., 2004) was used as a parent strain for all *B. subtilis* strains. A parent strain (Δ*rsbPQ* Δ*rsbTU rsbV-FLAG amyE::ctc-lacZ*) was generated for the screen of *rsbT*-independent *rsbU* variants. To generate a marked deletion of *rsbPQ,* the genomic regions 500 bp upstream and downstream of *rsbPQ* were amplified and ligated to a kanamycin resistance cassette using standard SOE PCR techniques. The linear DNA fragment was inserted into KC479 using standard techniques. Transformants were selected based on kanamycin resistance and confirmed by colony PCR and sequencing. A clean deletion of *rsbTU* was generated by generating a gBlock (IDT DNA Technologies) of regions corresponding to 500 bp upstream of *rsbT* and 500 bp downstream of *rsbU* ligated together. This fragment was introduced into the HindIII and EcoRI sites of pminiMAD2 (Arnaud et al., 2004). Standard genetic techniques were used to excise the chromosomal copy of *rsbTU*. Mutants were confirmed by colony PCR and sequencing of the genomic region. A single FLAG tag was introduced at the C-terminus of *rsbV* using the pminiMAD2 vector. Point mutations of the *rsbU* gene were similarly introduced using the pminiMAD2 vector and confirmed by colony PCR and sequencing.

**Table 1.**

| Alias | Species | Background | Construct/Genotype | Reference |
| --- | --- | --- | --- | --- |
| NB181 | *E. coli* | BL21(DE3)<br>rosetta pLysS | pET23a H6-Sumo SpoIIAA Rosetta2<br>plysS | Bradshaw and<br>Losick 2015 |
| NB182 | *E. coli* | BL21(DE3)<br>rosetta pLysS | pET23a H6-Sumo SpoIIAB<br>BL21(DE3) | Bradshaw and<br>Losick 2015 |
| NB781 | *E. coli* | BL21(DE3) | pET 6H-spoIIAA + spoIIAB in<br>BL21(DE3) | Levdikov et al.<br>2012 |
| NB1732 | *E. coli* | BL21(DE3) | BL21 (DE3) pET47b 6HIS-3C-SpoIIE 590-827 | Bradshaw et<br>al. 2017 |
| NB1937 | *E. coli* | BL21(DE3) | pET47b 6His-3C-spoIIE 590-827<br>V697A kan BL21 (DE3) | Bradshaw et<br>al. 2017 |
| NB1731 | *E. coli* | BL21(DE3) | BL21 (DE3) pET47b 6HIS-3C-SpoIIE 457-827 | Bradshaw et<br>al. 2017 |
| NB1878 | *E. coli* | BL21(DE3) | BL21 (DE3) pET47b 6HIS-3C-SpoIIE 457-827 V697A | Bradshaw et<br>al. 2017 |
| NB1833 | *E. coli* | BL21(DE3) | B. subtilis RsbT 3C 6His in pET47b<br>kan BL21(DE3) | This work |
| NB1834 | *E. coli* | BL21(DE3) | B. subtilis RsbU 3C 6His in pET47b<br>kan BL21(DE3) | This work |
| NB1835 | *E. coli* | BL21(DE3) | B. subtilis RsbV 3C 6His in pET47b<br>kan BL21(DE3) | This work |
| NB1836 | *E. coli* | BL21(DE3) | B. subtilis RsbW 3C 6His in pET47b<br>kan BL21(DE3) | This work |
| NB1925 | *E. coli* | BL21(DE3) | BL21 (DE3) pET47b 6His-3C-RsbVW kan | This work |
| KH170 | *B. subtilis* | PY79 | $\Delta rsbPQ \Delta rsbTU rsbV\text{-FLAG}$<br>$amyE::ctc\text{-lacZ}$ | This work |
| KH07 | *B. subtilis* | PY79 | $\Delta rsbPQ \Delta rsbTU rsbV\text{-FLAG}$<br>$amyE::ctc\text{-lacZ}$ c. pHB201-P(spank)-*rsbU* | This work |
| KH11 | *B. subtilis* | PY79 | $\Delta rsbPQ \Delta rsbTU rsbV\text{-FLAG}$<br>$amyE::ctc\text{-lacZ}$ c. pHB201-P(spank)-*rsbTU* | This work |
| KH109 | *B. subtilis* | PY79 | $\Delta rsbPQ \Delta rsbTU rsbV\text{-FLAG}$<br>$amyE::ctc\text{-lacZ}$ c. pHB201-P(spank)-*rsbU* M166V | This work |
| KH110 | *B. subtilis* | PY79 | $\Delta rsbPQ \Delta rsbTU rsbV\text{-FLAG}$<br>$amyE::ctc\text{-lacZ}$ c. pHB201-P(spank)-*rsbU* M166L | This work |
| KC479 | *B. subtilis* | PY79 | $amyE::ctc\text{-lacZ}$, *CmR* | Carniol et al<br>2004 |
| KH138 | *B. subtilis* | PY79 | $\Delta rsbPQ amyE::ctc\text{-lacZ}$, *rsbU*<br>M166L | This work |
| KH142 | *B. subtilis* | PY79 | $\Delta rsbPQ \Delta rsbU$ amyE::*ctc-lacZ* | This work |
| KH169 | *B. subtilis* | PY79 | $\Delta rsbPQ$ amyE::*ctc-lacZ* | This work |
| KH28 | *E. coli* | DH5α | pHB201-P(spank)-*rsbU* | This work |
| KH30 | *E. coli* | DH5α | pHB201-P(spank)-*rsbTU* | This work |
| KH130 | *E. coli* | DH5α | pHB201-P(spank)-*rsbU* M166V | This work |
| KH132 | *E. coli* | DH5α | pHB201-P(spank)-*rsbU* M166L | This work |
| KH173 | *E. coli* | DH5α | pminiMAD2-$\Delta rsbTU$ | This work |
| KH172 | *E. coli* | DH5α | pminiMAD2-$\Delta rsbU$ | This work |
| KH171 | *E. coli* | DH5α | pminiMAD2-*rsbU* M166L | This work |

An inducible expression vector (pHB201-P(spank)-*rsbU*) was generated to screen for *rsbT-*independent variants of *rsbU.* A gBlock (IDT DNA Technologies) was generated containing a super-folding *gfp* gene flanked by AgeI and NotI sites and introduced into plasmid pDR110 (Carniol et al., 2005) at the HindIII and SphI sites using standard genetic techniques. The fragment of the resulting plasmid containing the *lacI* gene and the inducible *sfgfp* gene were PCR amplified and inserted into vector pHB201 at the SacI and KpnI sites using standard genetic techniques to generate pHB201-P(spank)-*sfgfp*.

To generate expression constructs for *rsbU* and *rsbTU,* the appropriate genes were PCR amplified from genomic DNA from PY79 using a high-fidelity polymerase and introduced at the AgeI and NotI sites using standard genetic techniques.

To generate libraries of pHB201-P(spank)-*rsbU* containing mutagenized *rsbU,* the *rsbU* gene was PCR amplified from genomic DNA from PY79 using error-prone PCR, utilizing the natural error rate of Taq polymerase. Mutagenized *rsbU* was introduced at the AgeI and NotI sites using standard genetic techniques. Plasmids were introduced into the parent strain (Δ*rsbPQ* Δ*rsbTU rsbV-FLAG amyE::ctc-lacZ*) using standard *B. subtilis* transformation techniques.

### Screen

Six independent pools of mutagenized pHB201-P(spank)-*rsbU* were introduced separately into independent clones of the parent strain. Cultures were grown in selective media and plated on selective medium containing IPTG and X-gal for 2 days at 37C. Approximately 12,000 mutant strains were analyzed. Blue colonies were selected and phenotypes were confirmed by re-streaking on indicator medium before plasmids were miniprepped and sequenced. Mutations were re-introduced into pHB201-P(spank)-*rsbU* and introduced into fresh parent backgrounds to confirm that phenotypes were dependent on expression of mutated *rsbU*.

### Liquid based beta-galactosidase assays

Beta-galactosidase assays to measure SigB-reporter activity were adapted from previously described protocols (Harwood and Cutting, 2010). In brief, cells were grown to desired OD, spun down, and resuspended in Z-buffer (60 mM Na_2_HPO_4_ 7H_2_0, 40 mM NaH_2_PO_4_ H_2_0, 10 mM KCl, 1mM MgSO_4_, 50 mM 2-mercaptoethanol) and kept on ice. Cells were arrayed in a 96 well plate and the OD_600_ was measured to assess cell culture density. Cells were lysed in Z-buffer containing a final concentration of 10 mg/mL lysozyme for 30 minutes at 37°C. *o*-nitrophenyl-β-D-galactoside (ONPG) was dissolved in Z-buffer (4mg/mL) and added to lysed cells to a final concentration of 0.67 mg/mL). Absorbance at 420 nm (to measure ONPG cleavage) and 550 nm (to control for light scattering caused by cell debris) was measured over 40 minutes and the rate of LacZ production was calculated from the slope of the linear phase of A_420_ – 1.75*A_550_ (between 300-1500 seconds). Miller units were calculated using the formula that 1 Miller Unit = 1000*(A_420_ – (1.75*A_550_))/(t*v*OD_600_); t is time in minutes, and v is the volume of the reaction in milliliters.

## Acknowledgements

Richard Losick’s career of discovery from the soil bacterium *B. subtilis* inspired this study. Rich has been extremely helpful in writing this manuscript and launching the Bradshaw lab. This work was funded by startup funds to N. Bradshaw from Brandeis University and by National Institutes of Health Grant GM18568 to Richard Losick. The authors also thank Chris Miller, Lizbeth Hedstrom, Dorothee Kern, Rachelle Gaudet, Julia Kardon, and members of the Bradshaw lab for discussions and for critical reading of the manuscript.

---

**Figure 2– Supplement 1.**
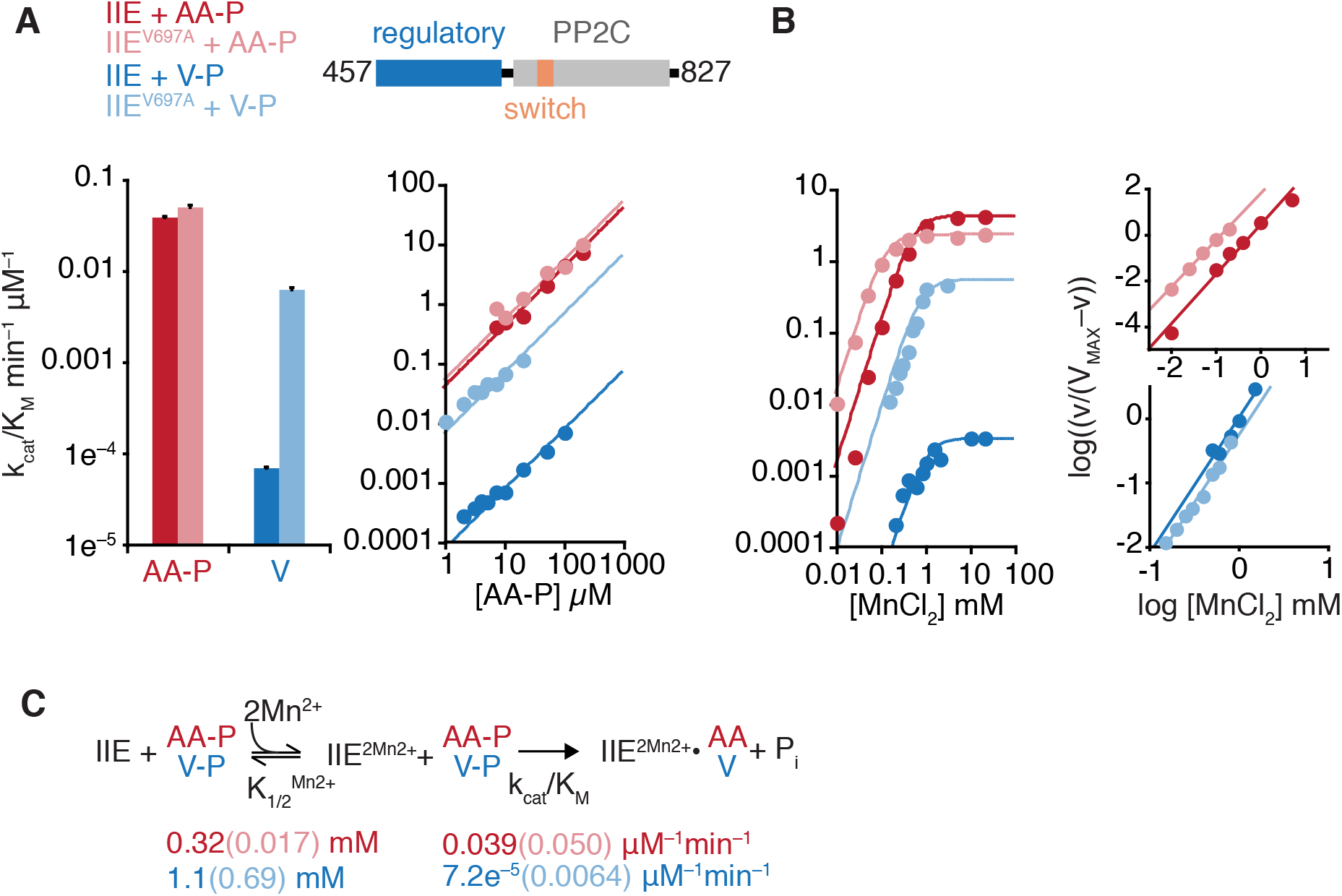
SpoIIE activity and specificity is influenced by the regulatory domain. **A** is a bar plot (left) summarizing the impact of the V697A substitution on the k_cat_/K_M_ of SpoIIE^457-827^. Data from SpoIIE^V697A^ are displayed in light colors. The bar plot is derived from the data displayed in the right graph. Data were fit to the equation observed rate = k_cat_*[SpoIIAA-P or SpoIIE]/(K_M_^MnCl2^+ [SpoIIAA-P or SpoIIE]). Error bars are the error of the fit. Reactions with SpoIIAA-P as substrate were performed as multiple turnover reactions with 0.1µM SpoIIE, 10mM MnCl_2_ and varying concentration of SpoIIAA-P. Reactions with RsbV-P as substrate were performed as single turnover reactions with 10mM MnCl_2_, trace RsbV-P and varying concentrations of SpoIIE (due to the slow rates of dephosphorylation). **B** is a plot of SpoIIE dephosphorylation of SpoIIAA-P (red) and RsbV-P (blue) as a function of MnCl_2_ concentration. Data are fit to a cooperative model using the equation k_obs_ = k_cat_*[MnCl_2_]^2^/(K_1/2_+ [MnCl_2_]^2^). Hill plots from the data are shown to the right and are fit to a linear equation. Data for SpoIIE^457-827^ and SpoIIE^457-827•V697A^ with SpoIIAA-P as substrate are from Bradshaw et. al 2017 and are replotted here as a reference. Reactions with RsbV-P as substrate included 50µM SpoIIE and trace RsbV-P. **C** is a summary of the reaction scheme for SpoIIE dephosphorylation with values determined for K_1/2_ for MnCl_2_, k_cat_/K_M_ below.

